## Supplementary Information S1-S6, Table S1, Figure S1-S6 for "Damage dynamics in single *E. coli* and the role of chance in the timing of cell death"

**Table of Contents**

**S1 Sample microscopy images and demonstration of the image analysis process**	**27**

**S2 Estimated of experimental noise**	**27**

**S3 Lifespan based on PI agrees with a different viability stain, AFH+TOPRO**	**28**

**S4 Best fit distribution functions to experimental damage distributions**	**29**

**S5 Shortening twilight in the E. coli dataset**	**31**

**S6 Analytical properties of the MP-SR model**	**32**

### S1 Sample microscopy images and demonstration of the image analysis process

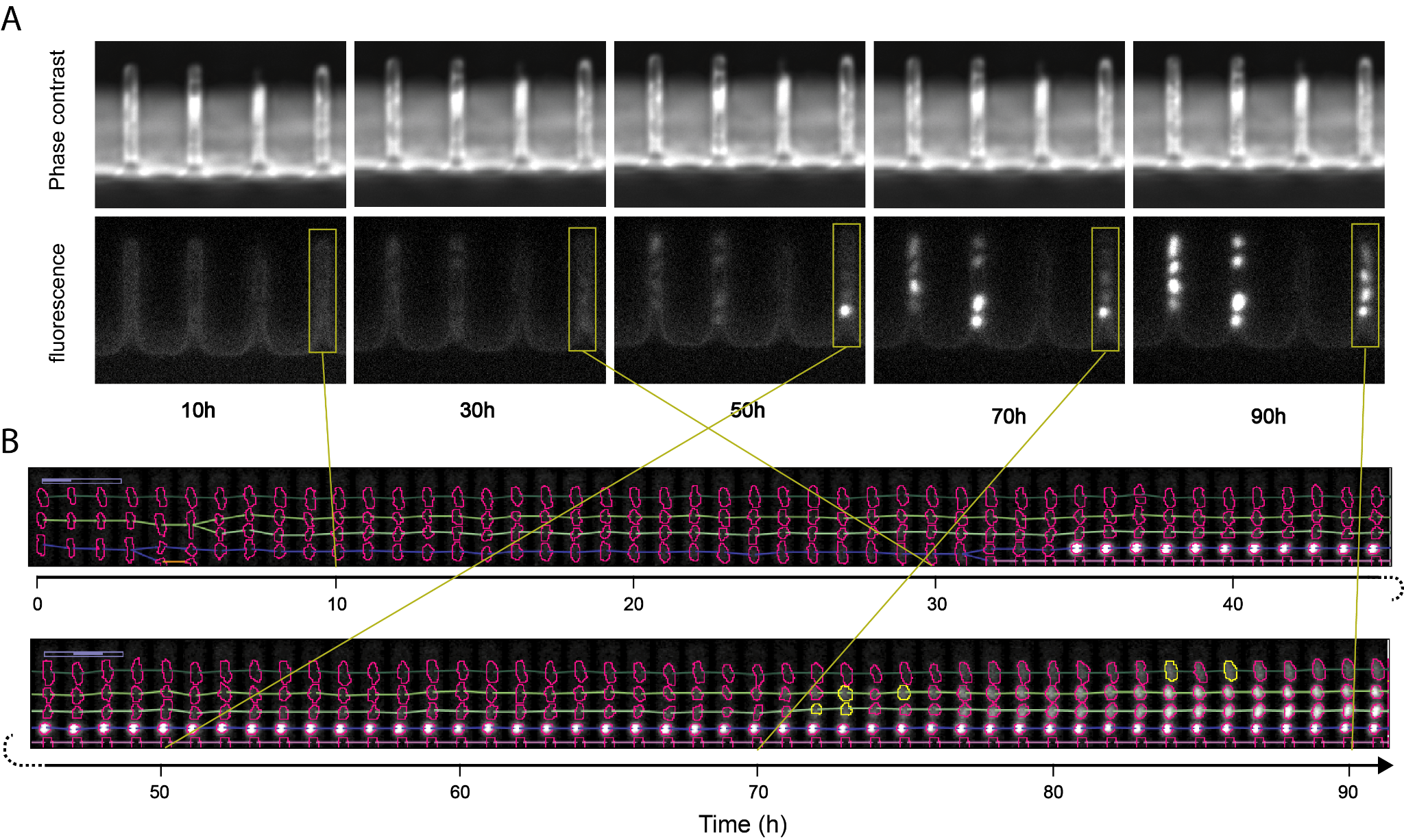

**Figure S1** Sample microscopy images (**A**) and demonstration of the image analysis process explained in Methods (**B**). Colored horizontal lines denote location and lineage tracking.

**Movie S1** Time-lapse movie of one sample imaging position. Each frame is a pseudo-color image, color-merged from CFP and PI fluorescence images.

### S2 Estimated of experimental noise

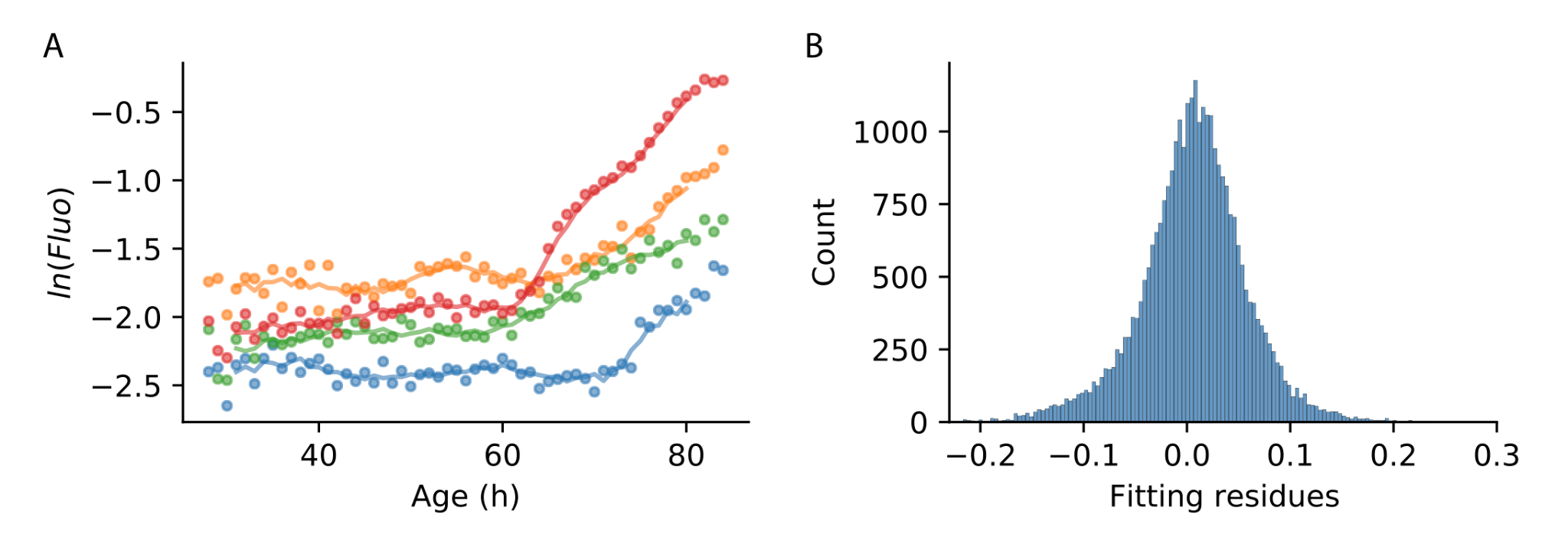

**Figure S2** Estimation of experimental noise. (**A**) Data derived from time-lapse microscopic images (dots) is modeled as a relatively slow moving signal (curves) and sequentially independent multiplicative experimental noise. The curves are moving averages of 7h windows with min and max values within each window removed. The estimate of experimental noise is the difference between these curves and the raw data in log scale, i.e. the fitting residues. (**B**) Histogram of the fitting residues. The mean of the distribution is approximately zero, 0.3%±0.4% per hour, and the standard deviation approximates the magnitude of experimental noise, about 5.5%. The left half of the distribution was used to estimate the magnitude of the experimental noise, because the distribution is skewed to the right due to the generally rising trends of PI signal.

### S3 Lifespan based on PI agrees with a different viability stain, AFH+TOPRO

We compared the present lifespan measurements using PI to viability using a different stain, AFH+TOPRO ^45^. This combined reagent only stains both damaged proteins (AFH) and DNA (TOPRO), and thus extends our main dye PI which stains DNA. We find good agreement between lifespan measured by PI in our microfluidic experiment and conventional single time-point (day 7) FACS assay of fraction surviving using AFH+TOPRO (Fig. S3)

We used PI in our experiments because of its previous validation in *E. coli* as a non-toxic dye to track cell viability longitudinally ^4,24^. Longitudinal experiments have different requirements than single-time-point experiments on cell viability - for example, uniformity of maximum intensity across cells might be more important in the latter. Longitudinal analysis allows detection of transient signals, and also captures the end-stages for each cell, and thus avoids some of the concerns inherent in single-time point studies.

In addition, our time-lapse microscopy experiments also capture not only the time course of the PI signal, but also the image of each cell throughout time (Figure S1, Movie S1). Thus we are able to confirm that the increases of PI signals are concentrated inside of the cells and centered around the nucleoids. These observations rule out the possibility of extracellular nucleic acids reported to occur in biofilms ^46^.

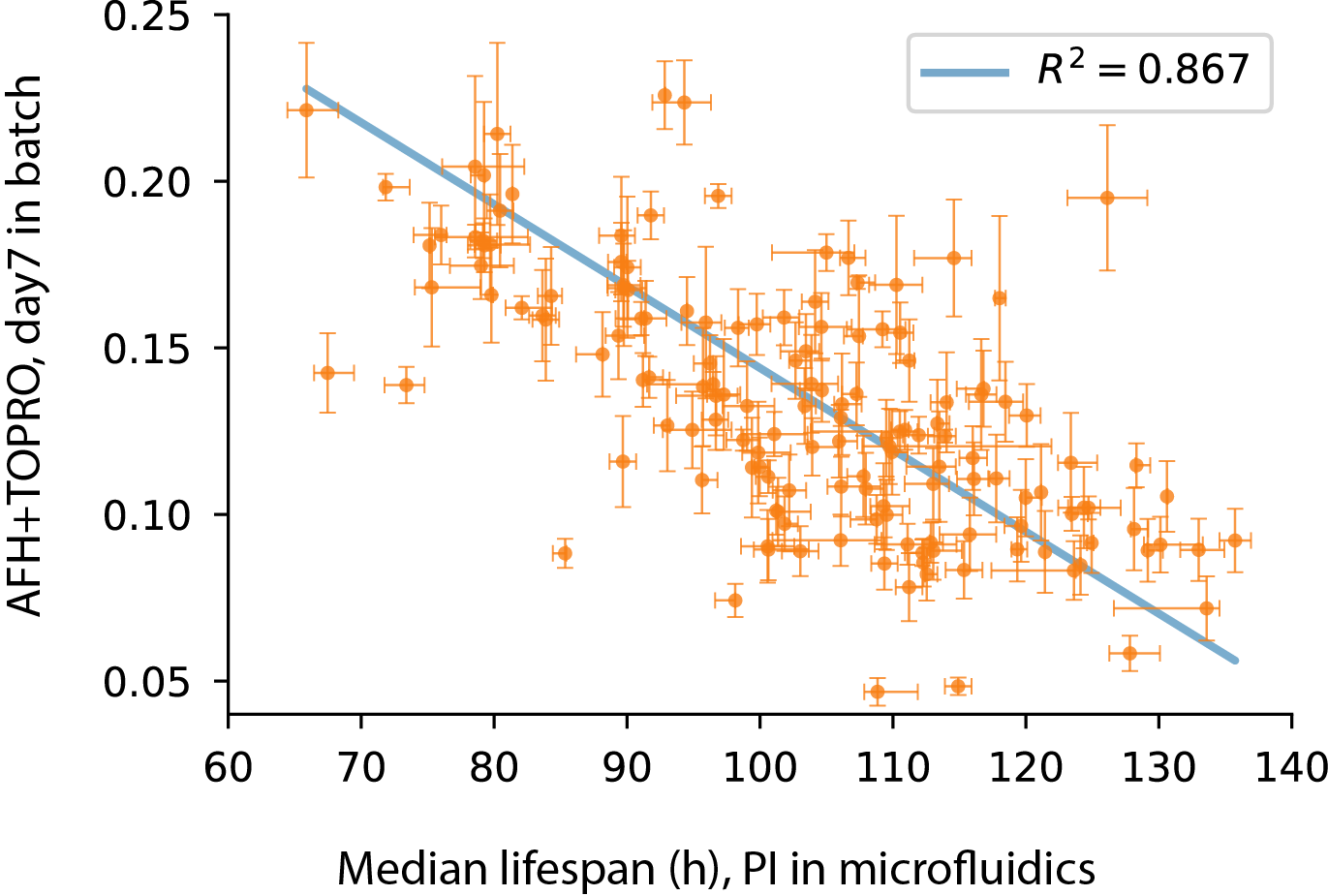

**Figure S3** Lifespan measurements using PI in microfluidic experiments correlate well with single-time point longevity measurements using AFH+TOPRO in batch culture. Each marker in orange corresponds to one *E. coli* strain (all based on MG1655 backgrounds). For each strain, we have performed microfluidic experiments ^4^ similar to those performed in the present study, in order to measure their survival curves. Plotted on the x-axis are median lifespan of these strains in microfluidic experiments. We also performed conventional batch-culture experiments to test cell viability, in conditions similar to those in microfluidics, and tested cell viability with AFH+TOPRO staining and FACS. Each train was tested three times. Plotted on the y-axis are the average fluorescence signal at day 7, normalized by cell size (estimated by forward scatter).

Fig. S3 include the lifespan measurements of the following strains, all based on MG1655 backgrounds: *wildtype, ∆agaA, ∆agaR, ∆agaV, ∆alsB, ∆appA, ∆appB, ∆appY, ∆aqpZ, ∆ascG, ∆bglF, ∆bipA, ∆caiB, ∆caiC, ∆caiT, ∆casE, ∆cbpM, ∆citT, ∆crcA, ∆cstA, ∆cynX, ∆dcp, ∆dcuA, ∆dnaT, ∆eptA, ∆eutQ, ∆fabR, ∆fixC, ∆fkpA, ∆flgD, ∆flu, ∆fruB, ∆fruR, ∆fsr, ∆glf, ∆glpX, ∆gltD, ∆gltK, ∆gltL, ∆gnsA, ∆gspF, ∆gspL, ∆gspM, ∆hslR, ∆hycA, ∆idnD, ∆intG, ∆ldcC, ∆lsrC, ∆lysC, ∆melA, ∆mhpB, ∆mprA, ∆mtlD, ∆nagK, ∆nagZ, ∆napC, ∆nirC, ∆norV, ∆norW, ∆nsrR, ∆ompL, ∆osmY, ∆paaC, ∆paoB, ∆pfkA, ∆pflC, ∆pflD, ∆pgi, ∆phnH, ∆ptsH, ∆rarD, ∆recF, ∆rph, ∆rsd, ∆sgcQ, ∆sspA, ∆ssuA, ∆ssuE, ∆syd, ∆tdcE, ∆thiQ, ∆treC, ∆tyrB, ∆ulaD, ∆uxaA, ∆uxaB, ∆uxaC, ∆uxuR, ∆xdhB, ∆xdhC, ∆yafV, ∆yafX, ∆yagH, ∆yagL, ∆yagQ, ∆yagS, ∆yajQ, ∆ybaJ, ∆ybbP, ∆ybdG, ∆ybdK, ∆ybiT, ∆ycaD, ∆ycaL, ∆ycfX, ∆ydcP, ∆ydgT, ∆ydjK, ∆ydjQ, ∆yecP, ∆yeeW, ∆yehA, ∆yfbM, ∆yfiD, ∆yfjO, ∆ygcQ, ∆ygfO, ∆yggE, ∆yggF, ∆ygjP, ∆yhbP, ∆yhbS, ∆yhbW, ∆yhdX, ∆yiaG, ∆yiaW, ∆yicH, ∆yicM*.

### S4 Best fit distribution functions to experimental damage distributions

In this section we provide details for the fits of the 15 distribution functions to the experimental *E. coli* damage distributions at different ages. We provide the KS statistics (Table S1) and p-values (Fig. S4), where a higher p-value (bluer colors) means a better fit. The distributions are ordered according to goodness of fit (average log p-value).

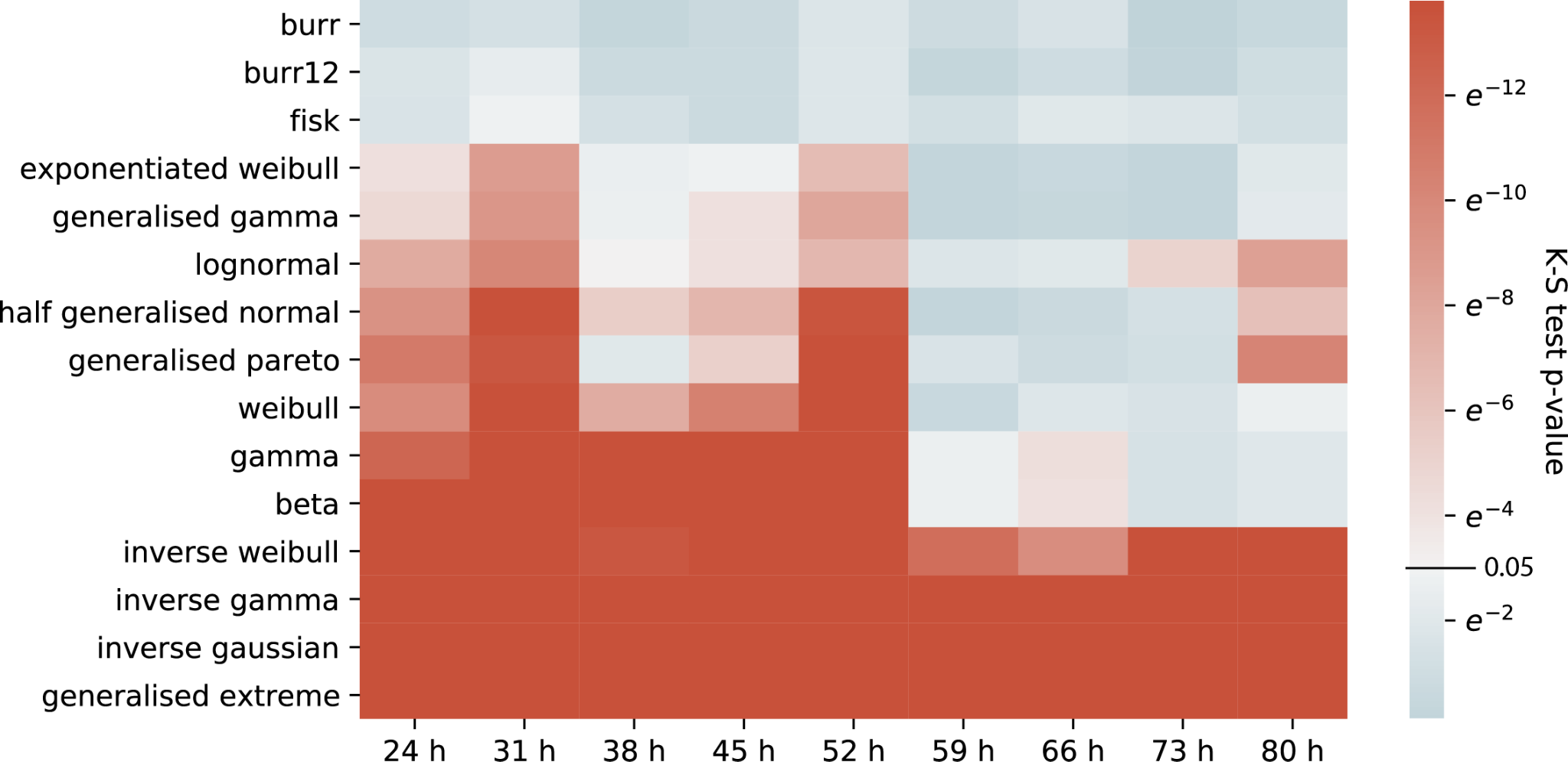

**Figure S4** KS test p-values for 15 distribution functions to the marginal damage distributions at different timepoints. Functions are ordered by goodness of fit (bluer is better).

| Model | K-S statistic | | | | | | | | |
| --- | --- | --- | --- | --- | --- | --- | --- | --- | --- |
|  | 24.5h | 31.5h | 38.5h | 45.5h | 52.5h | 59.5h | 66.5h | 73.5h | 80.5h |
| burr | 0.042 | 0.046 | 0.035 | 0.038 | 0.047 | 0.048 | 0.046 | 0.027 | 0.038 |
| burr12 | 0.051 | 0.057 | 0.041 | 0.038 | 0.048 | 0.038 | 0.040 | 0.029 | 0.046 |
| fisk | 0.049 | 0.061 | 0.048 | 0.038 | 0.048 | 0.052 | 0.051 | 0.051 | 0.048 |
| exponentiated weibull | 0.073 | 0.098 | 0.062 | 0.060 | 0.080 | 0.037 | 0.034 | 0.031 | 0.058 |
| generalized gamma | 0.076 | 0.101 | 0.063 | 0.069 | 0.088 | 0.036 | 0.033 | 0.032 | 0.059 |
| lognormal | 0.095 | 0.106 | 0.067 | 0.069 | 0.082 | 0.060 | 0.051 | 0.078 | 0.106 |
| half generalized normal | 0.104 | 0.129 | 0.085 | 0.087 | 0.112 | 0.036 | 0.036 | 0.045 | 0.093 |
| generalized pareto | 0.112 | 0.121 | 0.056 | 0.077 | 0.115 | 0.058 | 0.039 | 0.044 | 0.116 |
| weibull | 0.106 | 0.142 | 0.099 | 0.106 | 0.129 | 0.043 | 0.050 | 0.049 | 0.064 |
| gamma | 0.118 | 0.166 | 0.141 | 0.150 | 0.153 | 0.070 | 0.069 | 0.048 | 0.057 |
| beta | 0.127 | 0.166 | 0.140 | 0.149 | 0.156 | 0.070 | 0.068 | 0.048 | 0.057 |
| inverse weibull | 0.152 | 0.284 | 0.129 | 0.136 | 0.153 | 0.134 | 0.100 | 0.140 | 0.202 |
| inverse gamma | 0.214 | 0.461 | 0.185 | 0.239 | 0.253 | 0.223 | 0.135 | 0.195 | 0.281 |
| inverse gaussian | 0.220 | 0.784 | 0.201 | 0.280 | 0.279 | 0.326 | 0.152 | 0.216 | 0.316 |
| generalized extreme | 0.376 | 0.394 | 0.373 | 0.374 | 0.396 | 0.369 | 0.372 | 0.372 | 0.387 |

**Table S1** Kolmogorov-Smirnov (KS) test statistics for the 15 distribution functions compared to the marginal damage distributions at different timepoints.

### S5 Shortening twilight in the *E. coli* dataset

We follow the pioneering work of N. Stroustrup and colleagues and explore the question of twilight, the time between a measurable age-related phenotype to the time of death ^24^. Suppose there is an age-related phenotype that is equivalent to damage crossing a threshold X1. If we define twilight ^23^ as the remaining lifespan after the threshold is crossed, the question is whether twilight shortens or lengthens with the age at which the threshold is crossed.

The SR and MP-SR models predict that twilight shortens with age on average (Fig. S5ABC). Equivalently, the time to first cross X1, denoted $t_{1}$, is positively correlated with time of death $t_{d}$ (Fig 5SD), but with a correlation coefficient less than one (Fig. S5E). This prediction is borne out by the *E. coli* dataset (Fig. S5FG). A similar effect was observed in *C. elegans* ^24^.

The reason for shortening twilight in the model is that the damage production term $\eta t$rises with age. Individuals that cross X1 at early times have a low production term. It takes them longer (on average) to reach the death threshold than those crossing X1 at late times (Fig 1A). Thus there is a negative correlation between $t_{1}$ and remaining lifespan (Fig. S5EG).

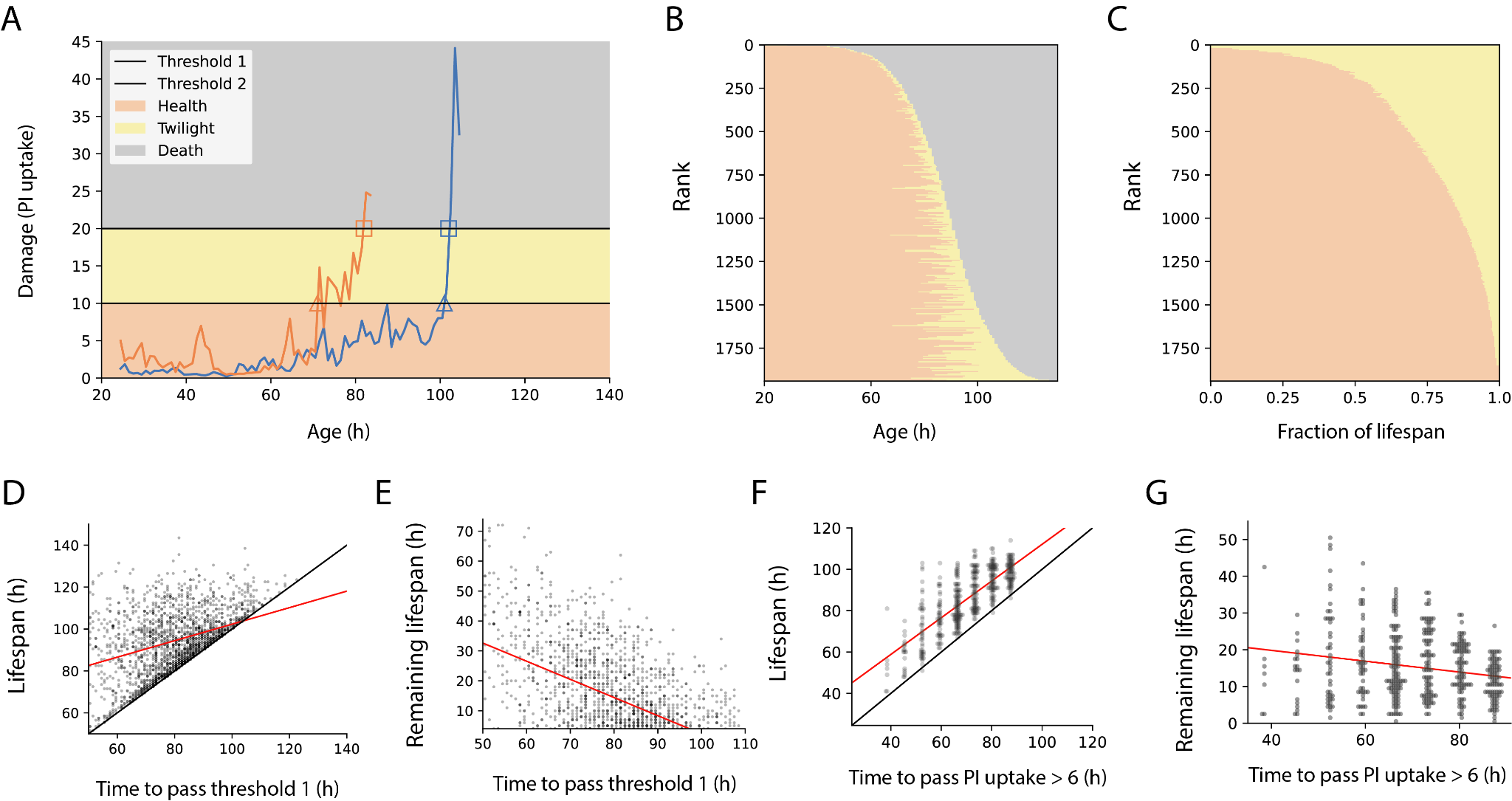

**Figure S5** *E. coli* shows shortening twilight at old age as redacted by the MP-SR model. (**A**) Examples of two MP-SR model simulation trajectories. Wildtype parameters were used. Thresholds are X1=10 for twilight onset and Xc=20 for death. Triangles and squares symbolize first-passage times ($t_{1}{,t}_{d})$ to cross the X1 and Xc respectively. (**B**) Durations of health ($t_{1}$, shown in red) and twilight ($t_{d}{-t}_{1}$, shown in yellow) for 2000 simulated cells ranked by lifespan. (**C**) Fraction of lifespan spent in health and twilight for the cells, ranked by fraction of time in health. (**D**) Time to first cross the two thresholds is correlated with slope less than 1 (Regression line y=0.39x+62.9) in MP-SR simulations. (**E**) Remaining lifespan drops with time to cross threshold 1 in MP-SR simulations. (**F**) *E. coli* lifespan versus the time to cross a damage threshold of normalized PI uptake rate=6 (Regression line is y=0.89x+23.3). (**G**) Remaining lifetime versus time to cross a damage threshold of normalized PI uptake rate =6.

### S6 Analytical properties of the MP-SR model

Here, we derive the risk of death of the MP-SR model analytically, using Kramer’s approximation ^20^.

The MP-SR model equation is:

$$dX/dt=\eta t - \beta e^{aX}/{(e}^{aX}+e^{a\kappa}) +\sqrt{2\sigma}\xi$$

One can write this in terms of a potential function $U(X,t)$ (Fig. 4C) :

$dX/dt=-\partial U(X,t)/\partial X +\sqrt{2\sigma}\xi$,

where the potential function is:

$U(X,t) = -\eta tX+\beta/a ln(e^{a\kappa}+e^{aX})$

We model mortality as the first time when $X>X_{c}$. Thus, death time is a first-passage time of the MP-SR model variable *X*. To estimate the risk of death, i.e. hazard rate, we apply the Kramer approximation ^21,22^ for the first passage time:

$$h(t)\approx\frac{\sqrt{U''(X_{0})U''(X_{C})}}{2\pi}e^{-\frac{U(X_{C})-U(X_{0})}{\sigma}}$$

Where $X_{0}$is the steady state of the system.

To arrive at Gompertz law, one needs $-\frac{U(X_{c})-U(X_{0})}{\sigma}$ to increase linearly with age. This is indeed the case:

$-\frac{U(X_{c})-U(X_{0})}{\sigma}=t\eta\sigma^{-1}(X_{c}-X_{0})+\beta a^{-1}\sigma^{-1}[z(X_{0})-z(X_{c})]$,

Where $z(x)=ln(e^{ax}+e^{a\kappa})$. If the quasi-steady-state $X_{0}$ was approximately constant and much smaller than Xc, as it is at young ages, one obtains the Gompertz hazard rate $h(t)\sim e^{\frac{\eta X_{c}}{\sigma}t}$ with the Gompertz slope $\eta\sigma^{-1}X_{c}$.

However, the quasi-steady state $X_{0}$ does increase with age, and rises more rapidly at late ages approaching Xc (Fig. 4B). Thus the hazard rate is only approximately Gompertzian, especially at late ages. If we use the quasi-steady state $X_{0}=ln(\frac{t\eta}{\beta-t\eta})/a$to calculate the hazard rate, we obtain a more complicated, non-Gompertzian formula, plotted in Fig. S6. This more realistic result shows late-age deceleration when compared with the Gompertz law. This deceleration is indeed observed experimentally for *E. coli* in similar conditions ^4^, as it is for other organisms ^20^. The deceleration also produces a survival curve that is better approximated by a Weibull survival curve than a Gompertz survival curve ^47^. The range over which the Gompertz law is accurate increases with $X_{c}/\kappa$.

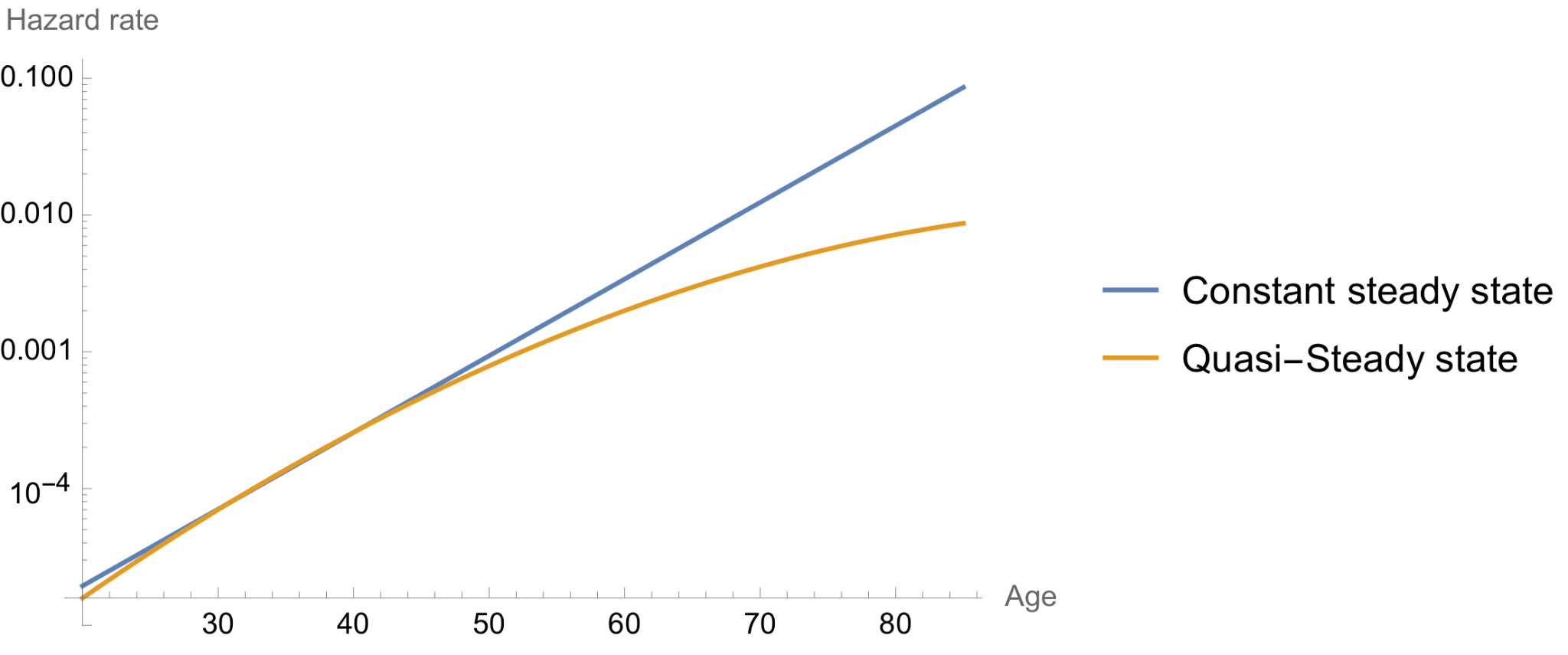

**Figure S6** Analytically calculated hazard rate of the MP-SR model using Kramer’s approximation. The two curves follow different assumptions: Blue curve shows Gompertz law, under a constant steady state assumption. The yellow curve uses a quasi-steady state that changes with age and shows late-age deceleration.
